# When rare meets common: Treatable genetic diseases are enriched in the general psychiatric population

**DOI:** 10.1101/2021.05.13.444051

**Authors:** Venuja Sriretnakumar, Ricardo Harripaul, James L. Kennedy, Joyce So

**Affiliations:** Campbell Family Mental Health Research Institute, Centre for Addiction and Mental Health, Toronto, Canada; Department of Laboratory Medicine and Pathobiology, University of Toronto, Toronto, Canada; Institute of Medical Science, University of Toronto, Toronto, Canada; Department of Psychiatry, University of Toronto, Toronto, Canada; Department of Medicine, University of Toronto, Toronto, Canada; Division of Medical Genetics, Departments of Medicine and Pediatrics, University of California, San Francisco, USA

**Keywords:** inborn errors of metabolism, channelopathies, schizophrenia, bipolar disorder, major depressive disorder, general anxiety disorder, obsessive compulsive disorder, psychiatric genetics

## Abstract

Mental illnesses are one of the biggest contributors to the global disease burden. Despite the increased recognition, diagnosis and ongoing research of mental health disorders, the etiology and underlying molecular mechanisms of these disorders are yet to be fully elucidated. Moreover, despite many treatment options available, a large subset of the psychiatric patient population is non-responsive to standard medications and therapies. There has not been a comprehensive study to date examining the burden and impact of treatable genetic disorders (TGDs) that can present with neuropsychiatric features in psychiatric patient populations. In this study, we test the hypothesis that TGDs that present with psychiatric symptoms are more prevalent within psychiatric patient populations compared to the general population by performing targeted next-generation sequencing (NGS) of 129 genes associated with 108 TGDs in a cohort of 2301 psychiatric patients. In total, 72 putative affected and 293 putative carriers for TGDs were identified, with known or likely pathogenic variants in 78 genes. Despite screening for only 108 genetic disorders, this study showed an approximately four-fold (4.13%) enrichment for genetic disorders within the psychiatric population relative to the estimated 1% cumulative prevalence of all single gene disorders globally. This strongly suggests that the prevalence of these, and most likely all, genetic diseases are greatly underestimated in psychiatric populations. Increasing awareness and ensuring accurate diagnosis of TGDs will open new avenues to targeted treatment for a subset of psychiatric patients.

## 1.2 Introduction

Mental illnesses are one of the biggest contributors to the global disease burden, with major depressive disorder (MDD) alone being the second highest contributor (Vigo et al., 2016). Despite the increased recognition, diagnosis and ongoing research of mental health disorders, the etiology and underlying molecular mechanisms of these disorders are yet to be fully elucidated. Moreover, despite many treatment options available, a large subset of the psychiatric patient population are non-responsive to standard medications and therapies (Bokma et al., 2019; Fabbri et al., 2019; Middleton et al., 2019; Mizuno et al., 2020; Sanches et al., 2019).

Decades of research has established that all of the common psychiatric illnesses - MDD, generalized anxiety disorder (GAD), schizophrenia spectrum disorders (SSD), bipolar disorder (BPD) and obsessive-compulsive disorder (OCD) - have contributing genetic factors (Gordovez et al., 2020; Liu et al., 2019; Meier et al., 2019; Ormel et al., 2019; Purty et al., 2019). However, there is a paucity of research and literature regarding the contribution of rare, highly penetrant genetic variants underlying psychiatric disorders. The limited studies have primarily focused on specific inborn errors of metabolism (IEMs), a subgroup of inherited genetic disorders wherein defects in proteins or enzymes along metabolic pathways result in toxic accumulations of substrates or metabolites (Bauer et al., 2013; Olivier Bonnot et al., 2019; Demily et al., 2017; Simons et al., 2017; Sriretnakumar et al., 2019; Trakadis et al., 2018), and chromosomal copy number variants (Charney et al., 2019; Cleynen et al., 2020; Rees et al., 2014; Sriretnakumar et al., 2019; Tansey et al., 2016). IEMs have been of particular interest due to the availability of targeted therapies and treatments for many of these disorders. (Saudubray et al., 2018; Waters et al., 2018). Although newborn screening and early presentation has led to many IEMs being primarily diagnosed in infancy and early childhood, many IEMs are now known to have late-onset presentations that are more prevalent than previously thought (Saudubray et al., 2009). Significantly, many late-onset IEMs present with psychiatric manifestations that can be indistinguishable from primary psychiatric disorders (Sedel et al., 2007; Sriretnakumar et al., 2019; Trakadis et al., 2018), and there have been numerous case reports of patients with late-onset IEMs who have been misdiagnosed as having a primary mental illness (Simons et al., 2017; Walterfang et al., 2013). Furthermore, heterozygote carriers for some autosomal recessive IEMs have been reported to present with psychiatric symptoms that can be ameliorated by treating the underlying genetic condition (Cocco et al., 2009; Tarnacka et al., 2009). Besides IEMs, there are other TGDs that can present with psychiatric phenotypes, including triplet repeat expansion disorders, neurocutaneous disorders and channelopathies (Cabal-Herrera et al., 2020; Kleopa, 2011; Northrup et al., 2018; Peng et al., 2018; Ratna et al., 2020). Examples of treatments for genetic disorders include specific drugs, such as miglustat for the treatment of Niemann-Pick disease type C (Patterson et al., 2020) and copper chelators for Wilson disease (Litwin et al., 2019), lifestyle modifications, such as avoidance of alcohol and fasting for acute intermittent porphyria (Fontanellas et al., 2016), dietary treatment, such as low-protein diet in urea cycle defects or phenylketonuria (Häberle et al., 2019), vitamin supplementation, such as folic acid, and vitamins B6 and B12 for homocystinuria (Jitpimolmard et al., 2020), and antiepileptic drugs for channelopathies, amongst many others (Baraban et al., 2013; Knupp et al., 2018; Pastor et al., 2018; Wolff et al., 2019). There is also some preliminary evidence of a role for variants associated with rare genetic disorders being associated with treatment non-responsiveness in psychiatric patients (Sriretnakumar et al., 2019), though further comprehensive study is needed.

There has not been a comprehensive study to date examining the burden and impact of TGDs in psychiatric populations. In this study, we test the hypothesis that TGDs that present with psychiatric symptoms are more prevalent within psychiatric patient populations compared to the general population by performing targeted next-generation sequencing (NGS) of 129 genes associated with 108 TGDs in a cohort of 2301 psychiatric patients.

## 1.3 Materials and Methods

### 1.3.1 Samples

A total of 2301 DNA samples from psychiatric patients were analyzed in this study. The patient cohort is a sub-sample retrieved from a larger sample set collected as part of the Individualized Medicine: Pharmacogenetic Assessment & Clinical Treatment (IMPACT) study at the Centre for Addiction and Mental Health (CAMH; Toronto, Canada). Sample characteristics are described elsewhere (Herbert et al., 2018; IMPACT, 2017). The demographic characteristics and psychiatric diagnoses of the current study cohort are provided in Table 4.1. Patient consent for genetic testing was obtained at the time of study recruitment, and research ethics board approval for this study was obtained through CAMH (Toronto, Canada).

**Table 4.1.** Demographic characteristics of the study sample (n=2 301).

|  | **SSD\*** | **BPD\*\*** | **OCD\*\*\*** | **MDD only** | **GAD only** | **MDD-GAD** | **Total** |
| --- | --- | --- | --- | --- | --- | --- | --- |
| **Number of Sample** | 436 | 556 | 375 | 179 | 302 | 453 | 2301 |
| **Male : Female %** | 308:128<br>71:29 | 177:379<br>32:68 | 161:214<br>43:57 | 46:133<br>26:74 | 105:197<br>35:65 | 123:330<br>27:73 | 920:1381<br>40:60 |
| **Ethnicity (%)** |  |  |  |  |  |  |  |
| **African** | 40 (9) | 6 (1) | 4 (1) | 3 (1) | 2 (1) | 5 (1) | 60 (3) |
| **Caucasian** | 298 (69) | 481 (86) | 320 (85) | 147 (82) | 260 (86) | 403 (89) | 1 909 (83) |
| **East Asian** | 18 (4) | 12 (2) | 8 (2) | 8 (5) | 7 (2) | 10 (2) | 63 (3) |
| **Hispanic** | 1 (0) | 5 (1) | 1 (0) | 4 (2) | 3 (1) | 4 (1) | 18 (1) |
| **Mixed Ethnicity** | 22 (5) | 13 (2) | 13 (4) | 7 (4) | 12 (4) | 11 (2) | 78 (3) |
| **Not Available** | 31 (7) | 33 (6) | 26 (7) | 10 (6) | 14 (4) | 15 (3) | 129 (6) |
| **Indigenous People (Canada)** | 5 (1) | 3 (1) | 0 (0) | 0 (0) | 2 (1) | 2 (1) | 12 (0) |
| **South Asian** | 21 (5) | 3 (1) | 3 (1) | 0 (0) | 2 (1) | 3 (1) | 32 (1) |

\*83 patients had a secondary psychiatric diagnosis. \*\*203 patients had a secondary psychiatric diagnosis. \*\*\*28 patients had a secondary psychiatric diagnosis. SSD, schizophrenia spectrum disorders; BPD, bipolar disorder; MDD, major depressive disorder; OCD, obsessive compulsive disorder; GAD, generalized anxiety disorder.

### 1.3.2 DNA Sequencing

NGS was performed for 129 genes associated with 108 TGDs that have been associated with psychiatric phenotypes (supplementary Table S4.1). Probes for targeted sequencing of the 129 genes were designed on the Agilent Technologies SureDesign online platform (https://earray.chem.agilent.com/suredesign/). DNA samples were purified using Agencourt AMPure XP (Beckman Coulter Life Sciences, Indianapolis, IN) and quantified using Qubit™ dsDNA BR Assay Kit (ThermoFisher Scientific Inc., Waltham, MA). Agilent Technologies (Santa Clara, CA) SureSelectXT protocol for 3µg of input DNA was followed for library preparation, hybridization, and capture. NGS was carried out on the NovaSeq SP flow cell (300 cycles) (Illumina Inc., San Diego, CA).

### 1.3.3 Bioinformatic Analyses

Sequence alignment, variant calling, and variant annotation were performed using in-house scripts on the CAMH Specialized Computing Cluster (see supplementary methods for more information). Variant classification was carried out based on American College of Medical Genetics guidelines (Richards et al., 2015) through the use of the Human Genetic Mutation Database (HGMD^®^) Professional 2020.1 (Stenson et al., 2017), ClinVar (Landrum et al., 2018) and Franklin by Genoox (https://franklin.genoox.com/clinical-db/home) platforms. Additional manual literature search was also conducted when discrepancies between the three platforms arose. See supplementary Table S4.3 for citations of all known pathogenic variants identified in this study sample.

### 1.3.4 Protein Modelling

Protein modelling was performed for likely pathogenic (LP) variants and variants of uncertain significance (VUSes) for disease genes where protein models were available from the Protein Data Bank (see supplementary Table S4.1 for further information). Protein modelling was carried out on PyMOL Molecular Graphics System Version 2.4 (Schrödinger LLC, New York City, NY). The most likely rotamer was chosen to depict the amino acid change of non-synonymous variants.

### 1.3.5 Statistical Analysis

Exact binomial test was used for statistical comparison of observed pathogenic variants versus expected variant frequencies. Expected variant frequencies were derived from established disease prevalence for autosomal dominant disorders and males affected with X-linked disorders, where available (supplementary Table S4.2). Expected carrier frequencies for autosomal recessive disorders and rates of females heterozygous for X-linked disorders were calculated based on disease prevalence and assuming Hardy-Weinberg equilibrium (HWE) when specific carrier rates were not available (supplementary Table S4.2). T-test was used to compare the mean number of medications in patients with and without variants of interest (i.e. LP or pathogenic variants). All statistical tests were performed using R version 3.6.0 (Team, 2017), all tests were 2-sided and all p-values were Bonferroni-corrected.

## 1.4 Results

### 1.4.1 Variants of Interest

Figure 4.1 summarizes the breakdown of all sequenced variants following bioinformatic analysis. All exons sequenced averaged 300X read depth per sample. A total of 1748 variants were identified following annotation and filtration. Of these, a total of 207 pathogenic and 215 LP variants were identified for further statistical comparison. All pathogenic and LP variants identified in the study population were either absent or found at very low frequencies in ethnicity-matched Genome Aggregation Database (gnomAD v2: http://gnomad.broadinstitute.org/) exome sub-populations (supplementary Table S4.3).

**Figure 4.1.**
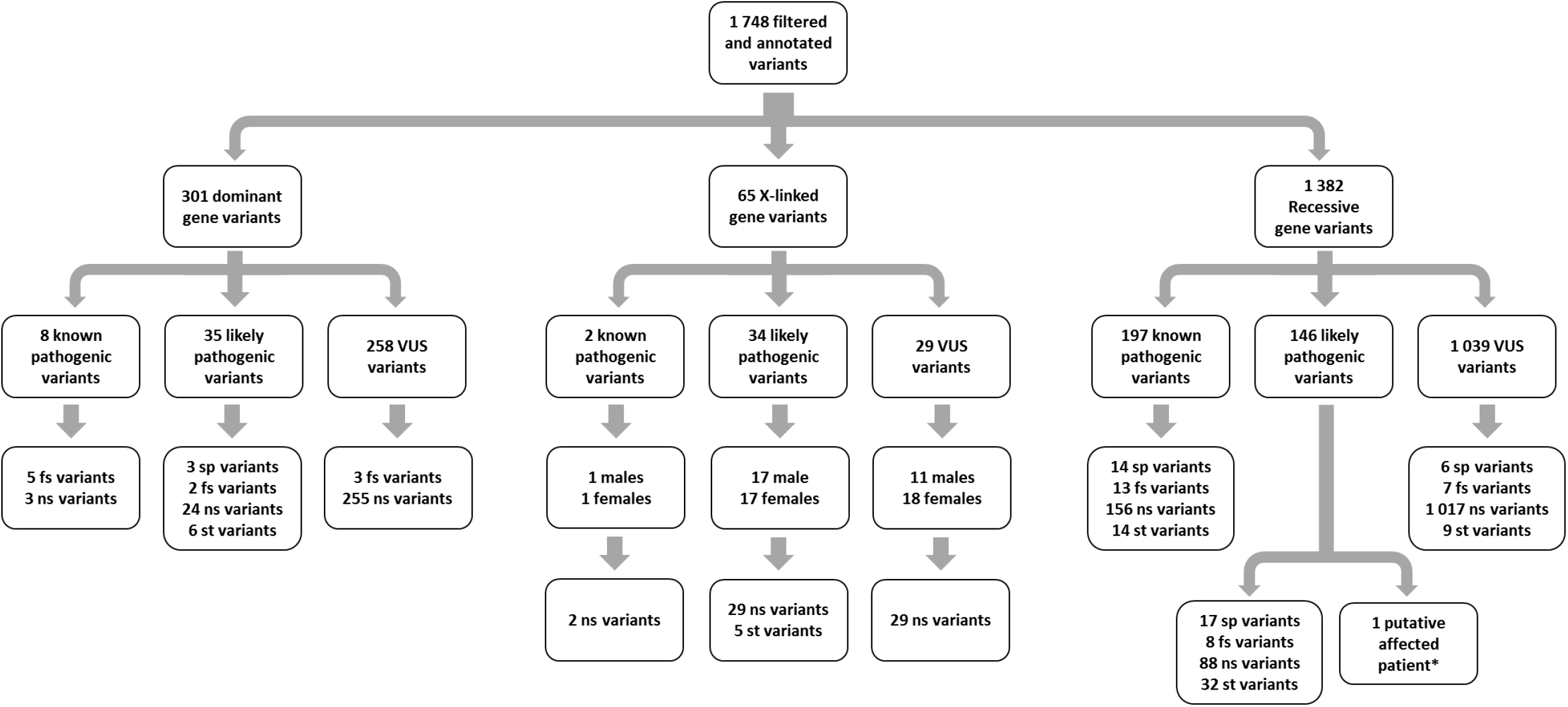
Diagram depicting the filtered and annotated variant breakdown in all 2301 SSD, BPD, OCD, MDD only, GAD only, and MDD-GAD samples. *One patient with 2 likely pathogenic variants in the same recessive gene. VUS, variant of uncertain significance; fs, frameshift; ns, nonsynonymous; sp, splice; st, stop gain/stop loss.

Table 4.2 shows the breakdown of patients identified with genetic variants of interest sorted by psychiatric diagnosis. Pathogenic or LP variants were identified in a total of 100 SSD, 64 BPD, 98 OCD, 22 MDD, 32 GAD and 49 MDD-GAD patients, with the highest frequency of pathogenic and LP variants identified in the OCD (39%) and SSD (28.2%) patient subsets.

**Table 4.2.**
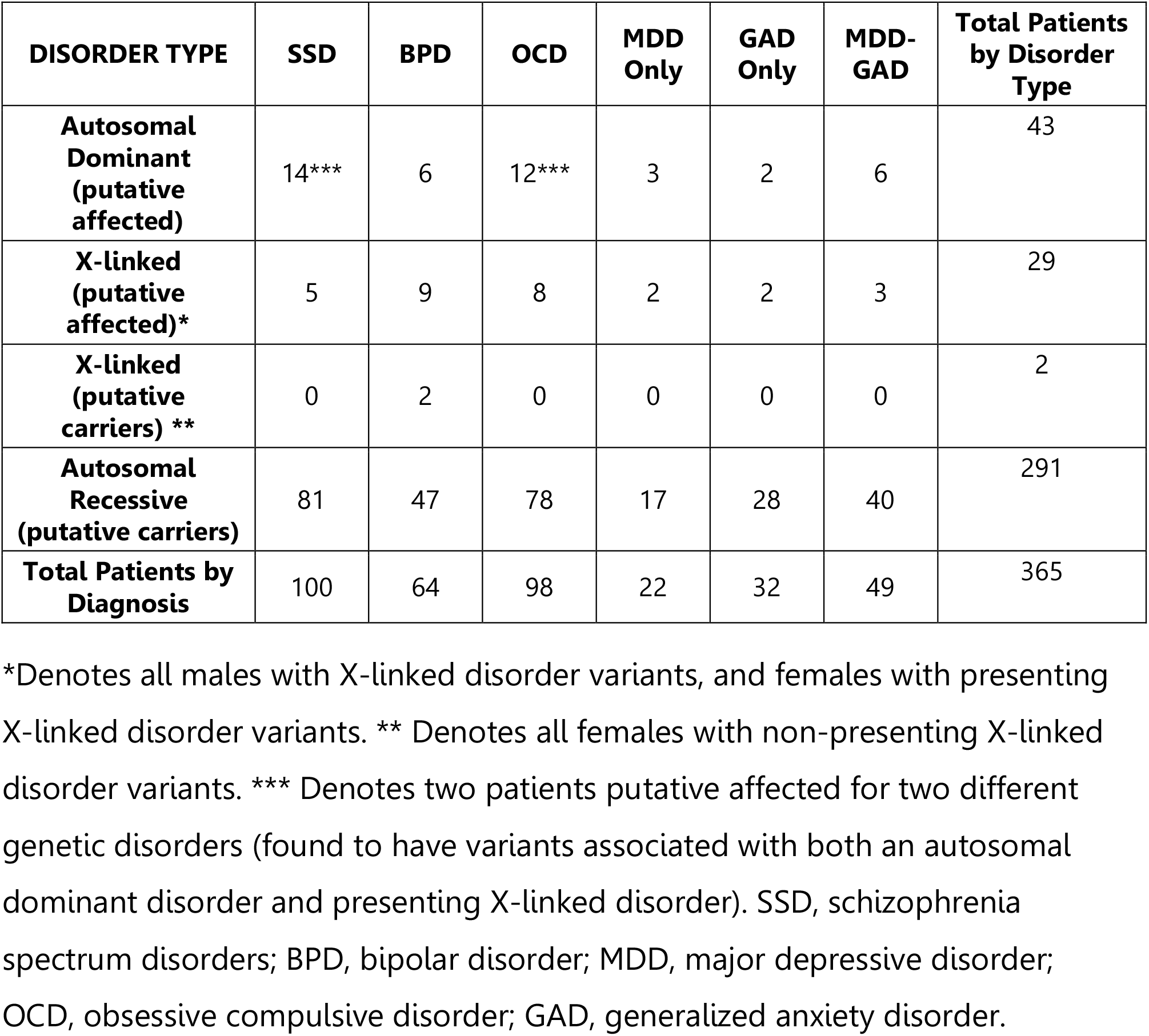
Number of patients with genetic variants of interest identified by psychiatric diagnosis.

There were also 22 patients identified to carry multiple variants within the same recessive disease-associated gene. Patient 6544 with OCD was identified to have two LP variants within the *ASL* gene. All other identified variants within this group were classified to be VUSes and thus were excluded from further statistical analysis.

### 1.4.2 Protein Modelling

Protein models were available to perform modelling of 40 LP variants and 29 VUSes (supplementary Tables S4.3 and S4.4). Of this, 33 LP missense variants resulted in changes to amino acid interactions within the protein, while seven LP variants did not result in any changes. However, two of the seven variants without protein interaction changes have previously known pathogenic variants at the exact same location and the remaining five variants are located in the same region as previously known pathogenic variants. For VUSes, 25 out of the 29 variants modeled resulted in significant amino acid interaction changes within the protein. Of the four variants that did not result in any protein changes, two variants are located in the same region as previously established pathogenic variants.

### 1.4.3 Prevalence of Treatable Genetic Disorder Variants within the Study Cohort

Patients with pathogenic or LP variants in genes associated with autosomal dominant disorders and male patients with pathogenic or LP variants in genes associated with X-linked disorder genes were considered to be putatively affected with the disorder. Those heterozygous for a pathogenic or LP variant in an autosomal recessive disorder gene and female patients heterozygous for a pathogenic or LP variant in an X-linked disorder gene were considered to be carriers for the disorder. The observed numbers of psychiatric patients who are putatively affected with and/or carriers for TGDs in comparison to expected disease prevalences and carrier frequencies are summarized in Table 4.3. The prevalences of autosomal dominant disorders related to pathogenic variants in *CACNA1A* (episodic ataxia type 2, familial hemiplegic migraine type 1, spinocerebellar ataxia type 6), *SCN1A* (epilepsy/febrile seizures, Dravet syndrome, familial hemiplegic migraine type 3), *SCN2A* (early infantile epileptic encephalopathy and/or benign familial infantile seizures), *SCN3A* (early infantile epileptic encephalopathy and/or benign familial infantile seizures), *COQ2* (susceptibility to multiple system atrophy),*HMBS* (acute intermittent porphyria; AIP), *PPOX* (porphyria variegate), *TTR* (hereditary amyloidosis), *NAGLU* (autosomal dominant Charcot-Marie-Tooth disease type 2V), and *GCH1* (dopa-responsive dystonia), and X-linked disorders related to pathogenic variants in *ABCD1* (X-linked adrenoleukodystrophy (X-ALD) and/or adrenomyeloneuropathy), *OTC* (ornithine transcarbamylase (OTC) deficiency), *PDHA1* (pyruvate dehydrogenase E1-alpha (PDHA1) deficiency) and *SLC6A8* (cerebral creatine deficiency syndrome 1) were significantly higher in the study cohort than expected based on general population prevalence, with the exception of OTC deficiency in females. There was an increased carrier rate for variants of interest in *COQ9* (biallelic mutations in which are associated with primary coenzyme Q10 (CoQ10) deficiency-5) and *GCDH* (biallelic mutations in which are associated with glutaric academia type 1 (GA1)) within the OCD subset of the study cohort relative to the general population.

**Table 4.3.**
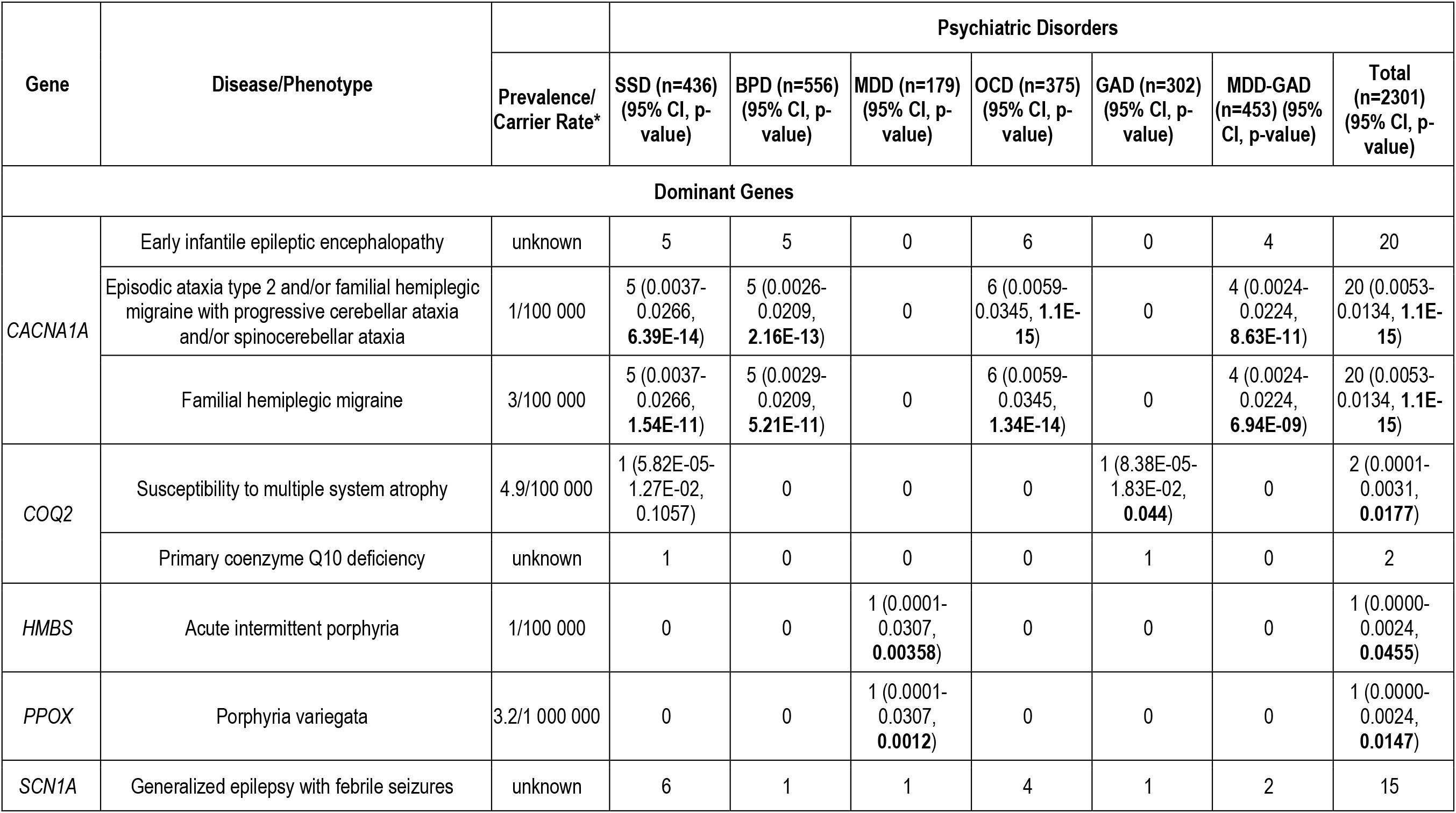
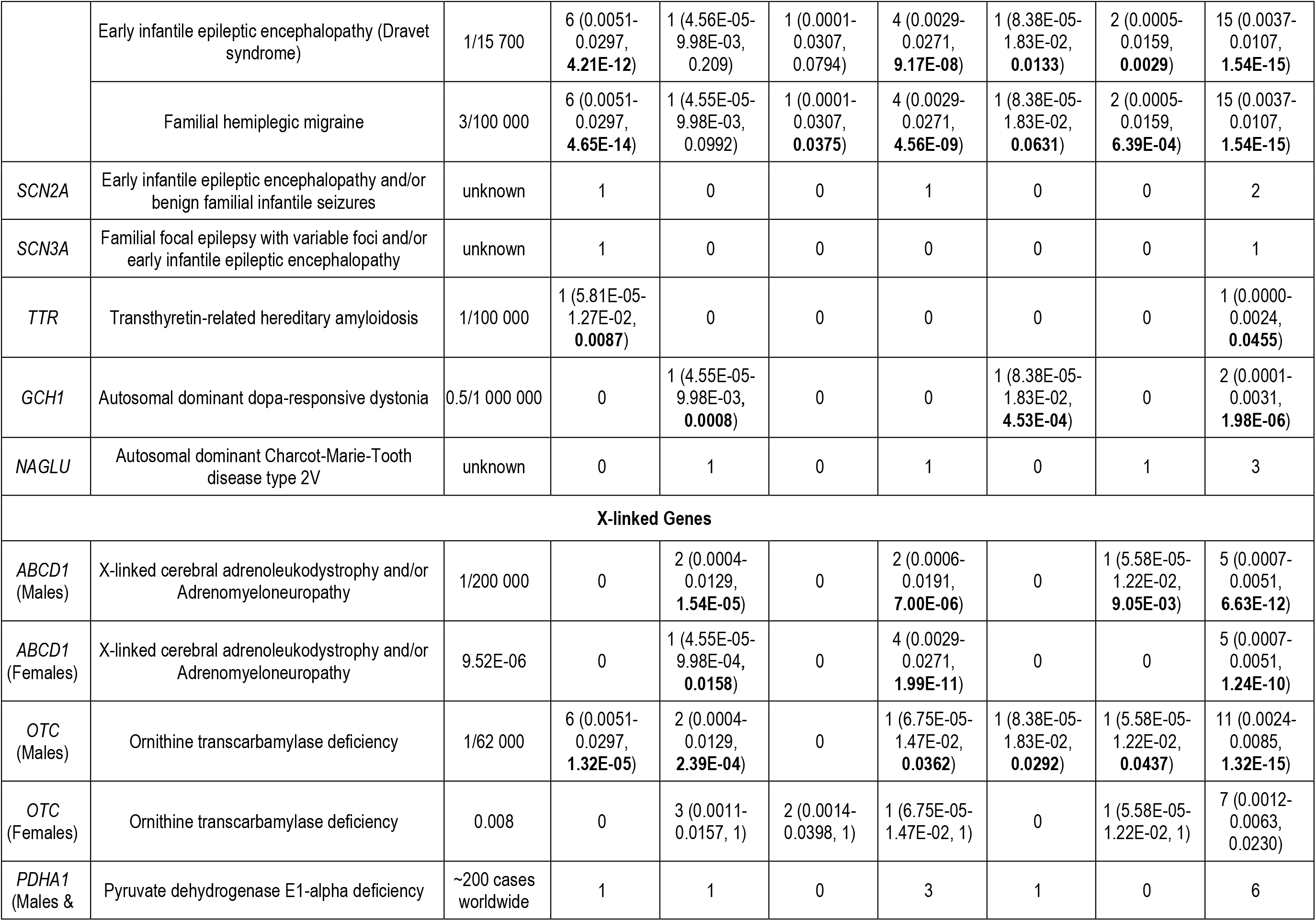
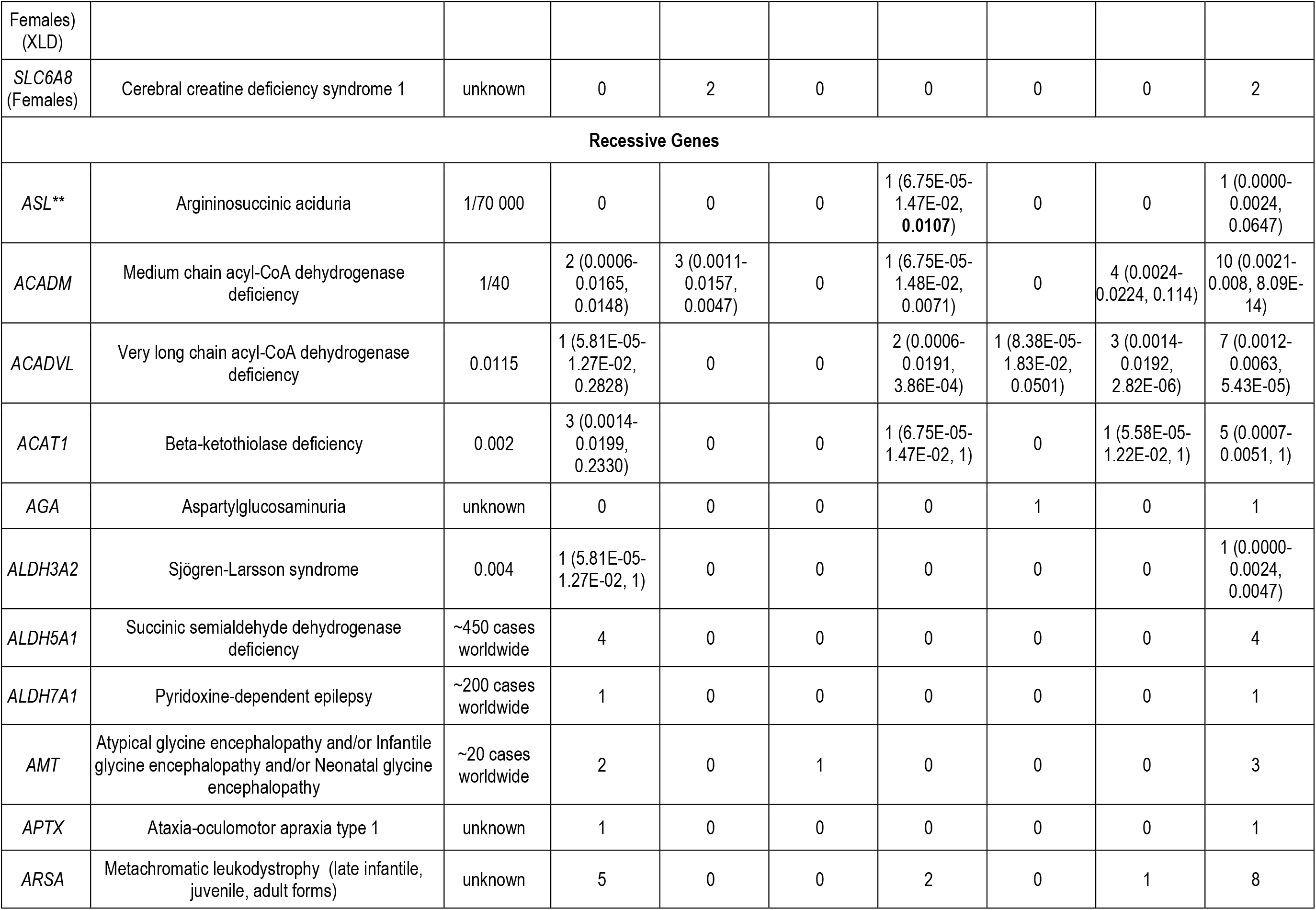
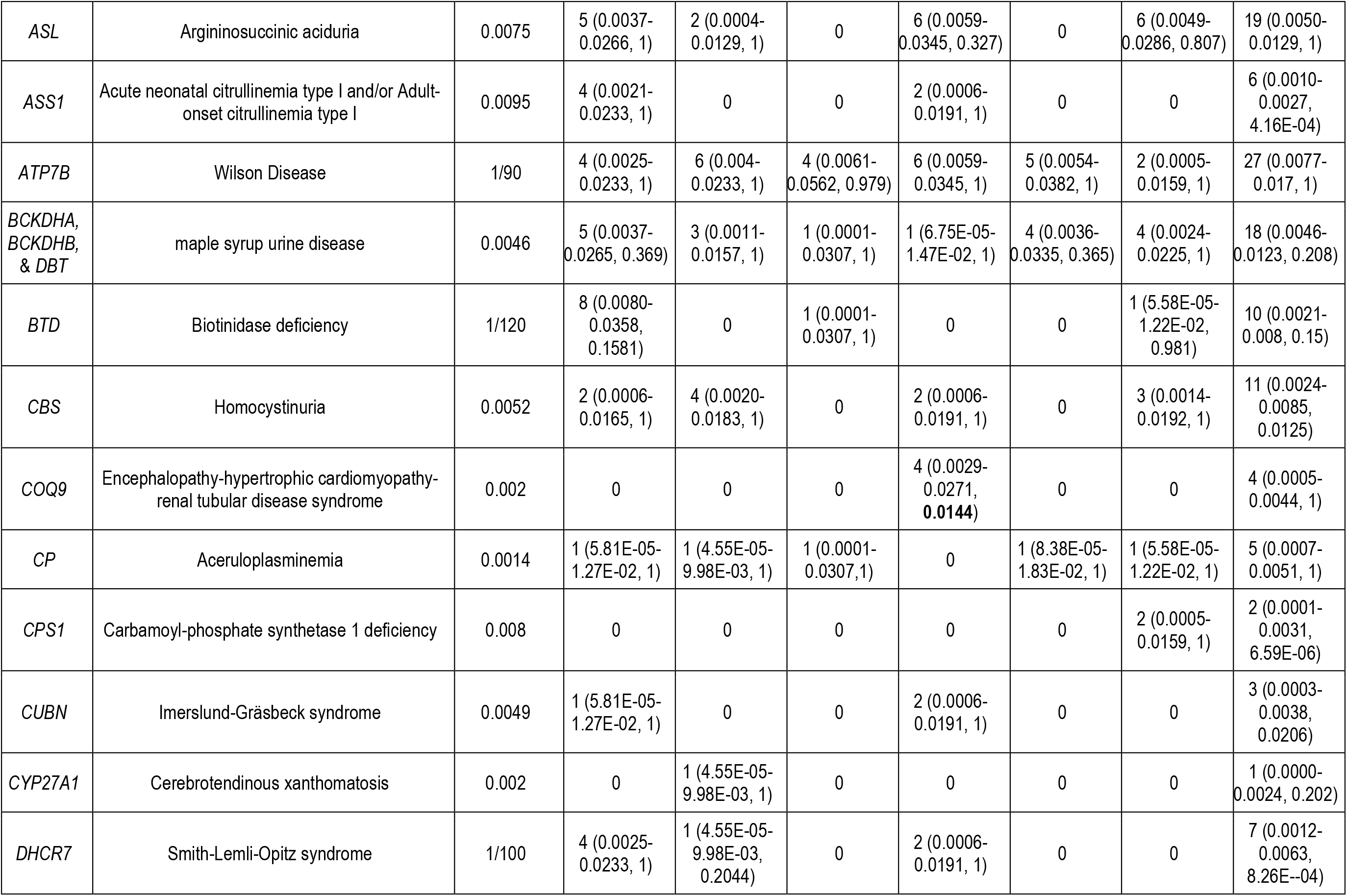
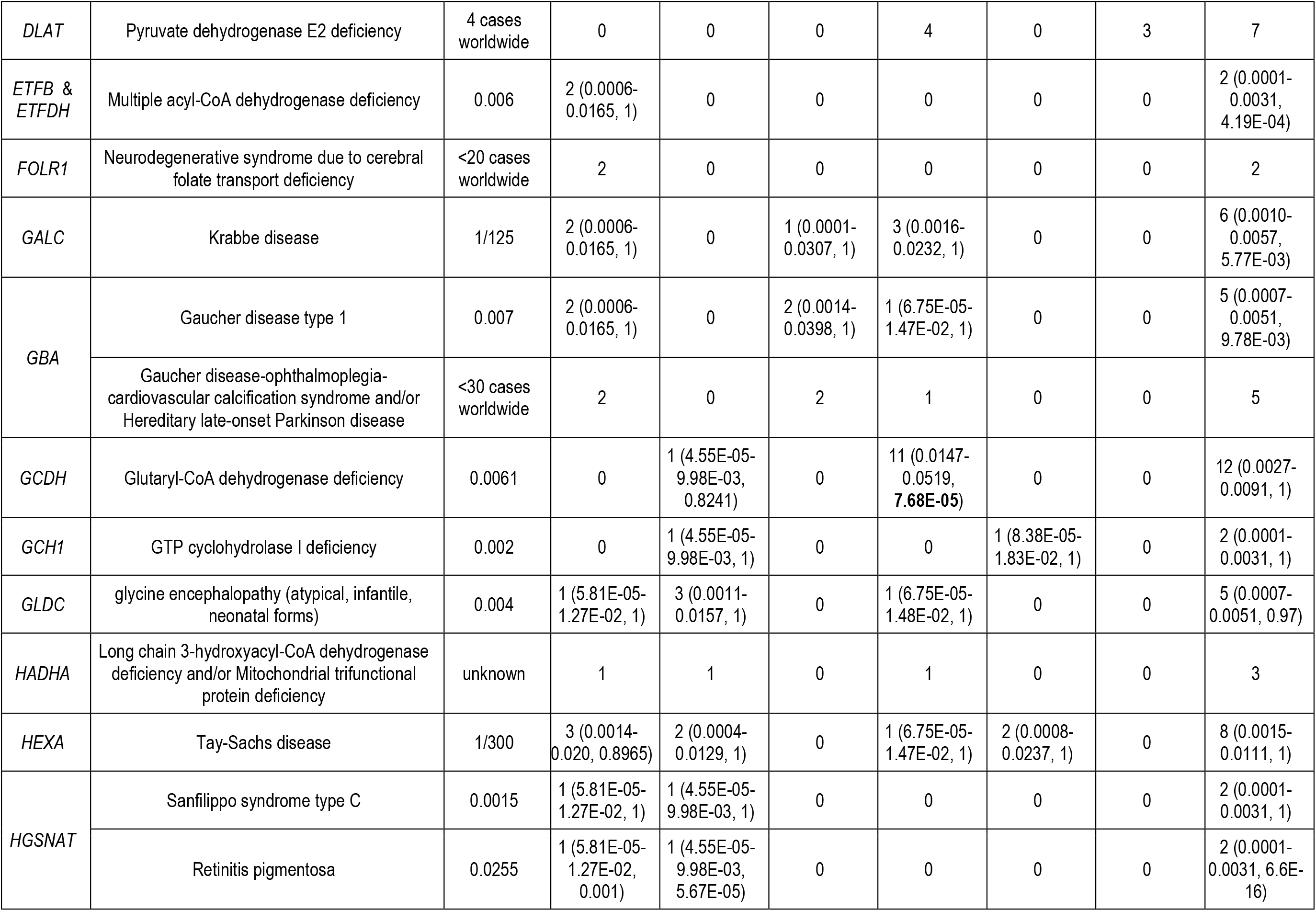
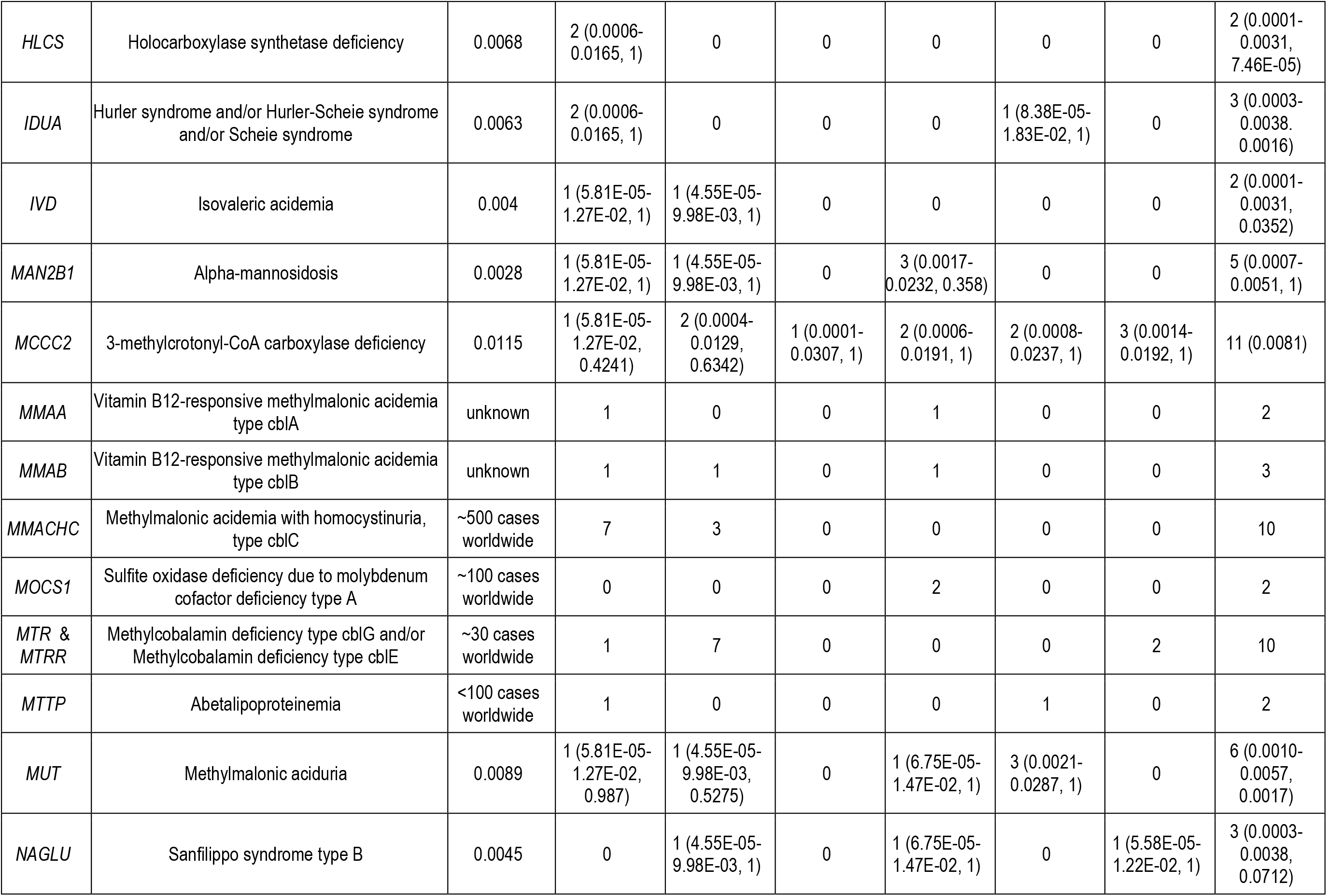
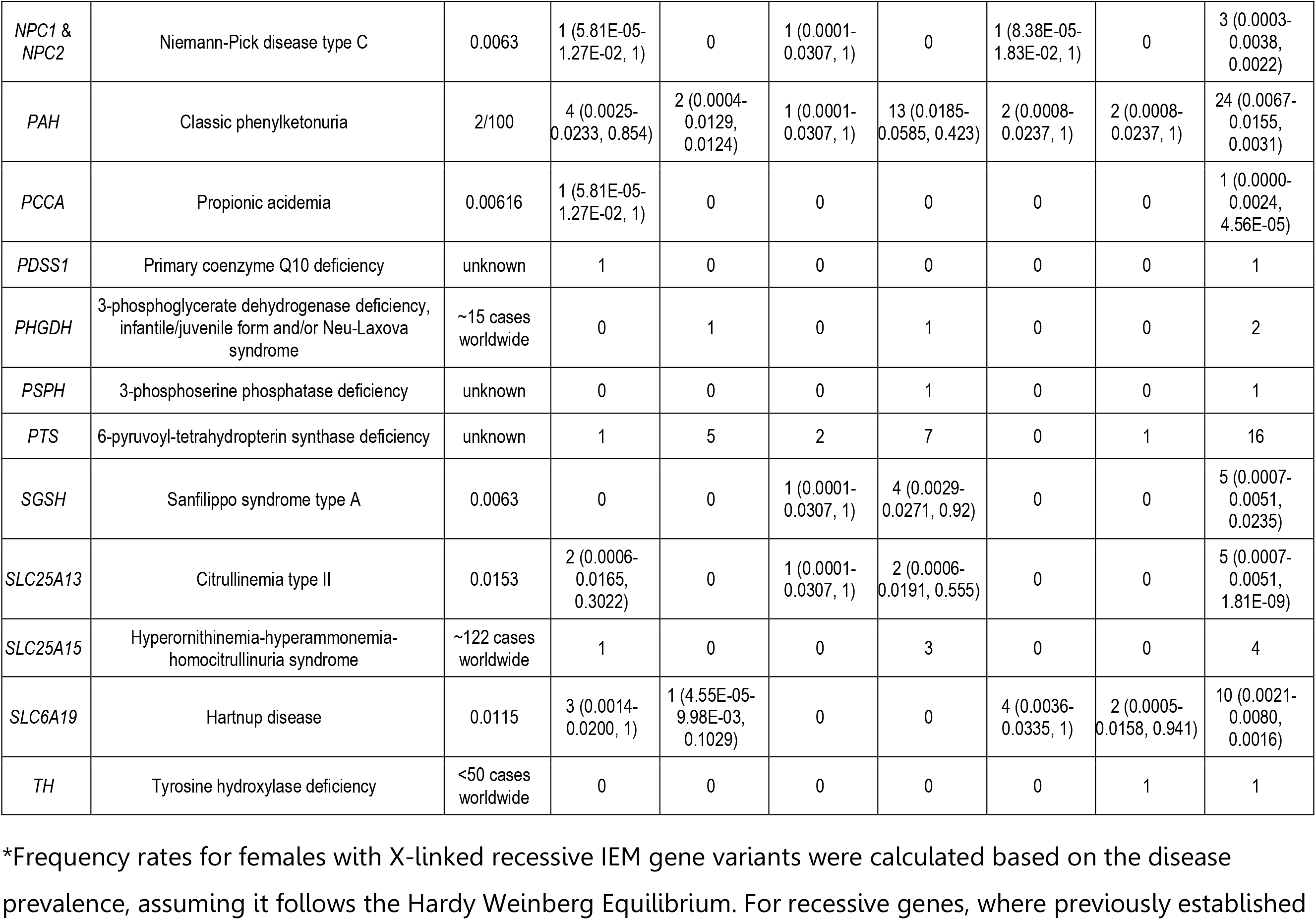
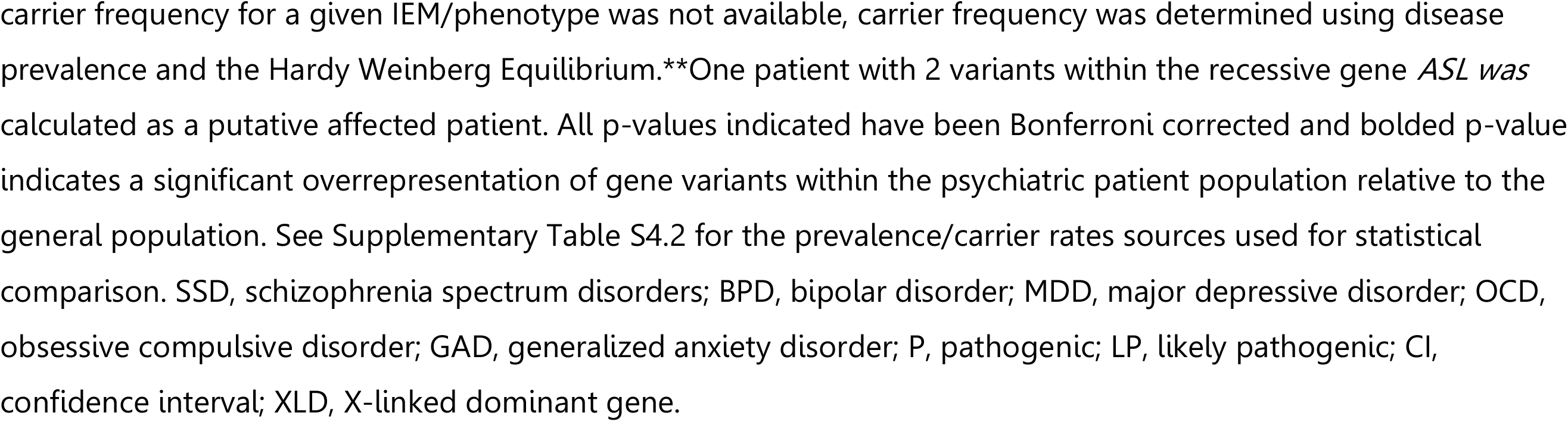
Prevalence of observed treatable IEM pathogenic variant frequencies in the psychiatric population compared to expected disease prevalence/carrier rates in the general population.

## 1.5 Discussion

This is the first study to directly investigate the prevalence of a comprehensive list of TGDs that are associated with psychiatric phenotypes in a large primary psychiatric cohort. Overall, 365 unique patients out of 2301 (15.86%) were identified to carry pathogenic or LP variants in TGD genes; 15 of these patients were found to have multiple variants of interests in genes associated with both dominant and recessive disorders. Of these, 62 (2.69%) individuals are putatively affected with a TGD, increasing to 72 (3.13%) with the inclusion of female heterozygous *OTC* and *ABCD1* pathogenic variant carriers, in whom variable penetrance and expressivity can affect the rate and severity of clinical presentation (Caldovic et al., 2015; Chongsrisawat et al., 2018; Engelen et al., 2014; Finsterer et al., 2013; Gyato et al., 2004; Schirinzi et al., 2019). A further 23 patients had VUSes that were predicted to cause structural effects on protein products, including 8 in autosomal dominant disorder genes, 2 in X-linked dominant disorder genes, and 13 patients with two variants within the same recessive disorder gene. Including these patients as potentially affected with TGDs would increase the proportion of putatively affected individuals within the psychiatric cohort to 4.13%. Taken together, the prevalence of TGDs in our study cohort is approximately three – and up to four - times the estimated 1% cumulative prevalence of all single gene disorders globally (Blencowe et al., 2018; WHO, 2020), despite only examining for 108 TGDs. This supports that the prevalence of all genetic disorders, including all single gene and chromosomal disorders, is likely much higher in psychiatric patient populations than previously thought. Additionally, 303 (13.17%) study patients are putative carriers for one or more of the screened autosomal or X-linked recessive disorders. This includes the discovery of pathogenic/LP variants in genes associated with very rare TGDs (e.g. primary coenzyme Q10 deficiency, cerebral creatine deficiency syndrome 1, among others), suggesting the potential enrichment of these extremely rare genetic diseases within psychiatric patient populations.

There are hundreds of IEMs with a collective prevalence estimated to be 50.9 per 100 000 live births (0.051%) and often overlooked is that many IEMs are treatable (Saudubray et al., 2018; Waters et al., 2018). In our previous pilot study (Sriretnakumar *et al*., 2019), we showed an enrichment of pathogenic variants in genes associated with four treatable IEMs (NPC, homocystinuria due to cystathionine beta-synthase deficiency, Wilson disease and AIP) in a cohort of SSD, BPD and MDD patients, with a putative affected rate of 0.34% with these four IEMs.. The current study identified 32 patients (1.39%) putatively affected with an IEM, representing a 27-fold enrichment of a select list of treatable IEMs within our psychiatric cohort relative to the collective prevalence of all IEMs. Major groups of enriched IEMs identified in this study include urea cycle disorders (n=49 affected patients), peroxisomal disorders (n=10) and porphyrias (n=2). Many of these disorders are highly treatable or manageable through diet, medications and lifestyle modifications (Diaz et al., 2013; Fontanellas et al., 2016; Häberle et al., 2019; Jiang et al., 2018; Mew et al., 2017; Morita, 2019; Pischik et al., 2015; S Grewal et al., 2016).

The enrichment of AIP – an autosomal dominant IEM - within the psychiatric population was replicated in this study from our previous study. This finding is in line with numerous studies in the literature demonstrating a strong association between AIP and psychiatric phenotypes (Duque-Serrano et al., 2018). Specifically, one study found a 20-fold increased incidence of AIP in hospitalized psychiatric patients in comparison to the general population, while another study showed AIP patients to be at a four-fold increased risk for being diagnosed with SSD and BPD, and, significantly, their first-degree relatives were found to have a two-fold increased risk for developing SSD and BPD (Cederlöf et al., 2015; Duque-Serrano et al., 2018; Tishler et al., 1985). Treatment options for the porphyrias include preventing or minimizing acute porphyric crises through avoidance of precipitating factors (e.g. alcohol, smoking, stress, certain medications), which can prevent the onset of psychiatric symptomatology altogether, as well as carbohydrate loading, electrolyte infusions and heme therapy to abrogate acute attacks (Pischik et al., 2015). There are also emerging novel treatments for AIP, including enzyme, gene and messenger RNA-based therapies (Fontanellas et al., 2016; Jiang et al., 2018; Parra-Guillen et al., 2020).

Urea cycle and peroxisomal disorders, such as the X-linked disorders OTC deficiency and X-ALD, have also been shown to present with psychiatric symptoms akin to SSD, BPD, attention deficit hyperactivity disorder (ADHD) and depression (Kitchin et al., 1987; Shamim et al., 2017; Stepien et al., 2019). OTC deficiency can present in infancy or childhood with seizures, neurodevelopmental impairment and ADHD (Lichter-Konecki et al., 2016). Although OTC deficiency is classically considered an X-linked recessive disorder, wherein males are affected with more severe phenotypes, it is well-known that females heterozygous for pathogenic *OTC* variants are also symptomatic, especially for neuropsychiatric phenotypes with a later onset (Lichter-Konecki et al., 2016; Lipskind et al., 2011; Niwinski et al., 2020; Pridmore et al., 1995). Interestingly, there was no enrichment of female carriers for OTC deficiency within the psychiatric cohort, though this may be confounded by the lack of an established carrier rate with which to accurately compare. Both males and females in the study cohort were enriched for pathogenic variants in *ABCD1*, the gene associated with X-ALD, in line with the reported 39% of X-ALD patients who present with one or more psychiatric symptoms, including, most notably, 17% of patients reported to present with exclusively psychiatric manifestations (Kitchin et al., 1987).

Taken together, the high rate of IEMs found in the study cohort suggests a very strong association between IEMs and psychiatric disorders. Whether the IEMs are causally related or modify existing psychiatric illnesses already present within a given patient is yet to be fully explored. Nevertheless, accurate genetic diagnosis of this subset of psychiatric patients with underlying IEMs is critical to allow for targeted therapy for the underlying disorder. The efficacy of targeted IEM therapies on psychiatric phenotypes specifically requires systematic study. There is clear precedent for positive effects on neuropsychiatric phenotypes from treatment for some common IEMs, such as phenylketonuria and OTC deficiency (Ashe et al., 2019; Gyato et al., 2004; Niwinski et al., 2020).

In this study, carriers for Wilson disease, metachromatic leukodystrophy, NPC and very long chain acyl-CoA dehydrogenase deficiency (VLCADD) were identified, amongst many other screened disorders. Heterozygous carriers of autosomal recessive disorders have been shown in previous studies to manifest with clinical findings, including psychiatric phenotypes. There are reports of heterozygous *ATP7B* mutation carriers presenting with neuropsychiatric features that have been alleviated through copper chelation, a standard treatment for full-fledged Wilson disease (Cocco et al., 2009; Tarnacka et al., 2009). Carriers for metachromatic leukodystrophy have been shown to have low arylsulfatase A (ARSA) activity, which has been strongly associated with neuropsychiatric features, including SSD and psychosis; specifically, heterozygous carriers for the ARSA I179S variant present with a psychiatric onset of metachromatic leukodystrophy which can mirror SSD, in addition to presenting with dementia and neurological phenotypes (e.g. paraparesis) (Demily et al., 2014; Ługowska et al., 2005; Marcão et al., 2003). Multiple studies have shown an enrichment for NPC carriers within psychiatric populations (Bauer et al., 2013; Maubert et al., 2013, 2015; Sriretnakumar et al., 2019). One patient with a heterozygous mutation in *ACADVL* was found to exhibit rhabdomyolysis, a clinical presentation of VLCADD due to biallelic *ACADVL* mutations (Hisahara et al., 2015). Patients both affected with and carriers for aspartylglucosaminuria have been found to have similar dysmorphic facial features, though no carriers for either disorder have been reported to have psychiatric phenotypes (Arvio et al., 2004). Although it cannot be excluded that a second mutation has been missed based on molecular testing limitations in these patients, it is worthwhile considering the possibility of manifesting carriers, which may pave the way for targeted therapies, as seen in the Wilson disease carrier examples. Future large studies to phenotype and potentially further genotype (e.g. with in-depth search for non-exonic variants) carriers for genetic disorders in psychiatric populations should be considered.

Interestingly, the highest proportion (39%) of pathogenic and LP variants in the genes studied was found in the OCD subgroup relative to the other psychiatric subgroups in the study cohort, with 20 of 375 OCD patients (5.3%) putatively affected and 78 (20.8%) carriers for recessive disorders. One potential explanation for this is that OCD has one of the most rigorous diagnostic criteria and, thus, the OCD patients within our study sample could have been enriched for more severe presentations of the illness relative to the other psychiatric sub-groups (Association, 2013). This is in line with our previous studies, in which we have shown that psychiatric patients enriched for genetic disease variants present with more severe psychiatric symptomatology in comparison to patients without these variants (Sriretnakumar et al., 2019).

Amongst the genes associated with treatable autosomal recessive disorders screened in this study cohort, there was an enrichment of heterozygous pathogenic/LP variants in *COQ9* and *GCDH* within the OCD subset of patients relative to the general population. Biallelic mutations in *COQ9* result in highly heterogeneous primary CoQ10 deficiency, features of which include cognitive deficits, intellectual disability, neuropathy, ataxia and seizures (Adam et al., 1993-2020). Interestingly, deficiency of CoQ10 is hypothesized to contribute to the well-recognized mitochondrial dysfunction seen in neuropsychiatric illnesses, such as SSD, BPD, MDD, Huntington disease and Parkinson’s disease (Maguire et al., 2018). CoQ10, an antioxidant, is found to be depleted in patients with neuropsychiatric disorders, and subsequent CoQ10 supplementation has been shown to have antidepressant effects and may slow progression of symptoms in Parkinson’s disease (Maguire et al., 2018; Morris et al., 2013). To date, no associations between *COQ9* and OCD have been found; however, OCD has been extensively associated with mitochondrial dysfunction and oxidative stress, the same pathways implicated in CoQ10 deficiency (Maia et al., 2019). Specifically, most OCD patients are found to have a dysregulated oxidative profile, wherein the elevated oxidative stress is insufficiently buffered by the antioxidant systems (Maia et al., 2019). Our study suggests a potential association between *COQ9* and OCD, which could potentially be mediated through cellular oxidative stress from mitochondrial dysfunction. Further study is warranted, particularly given the possibility of easy and accessible treatment with antioxidants, whether by dietary management or supplementation.

Biallelic mutations in *GCDH* result in glutaryl-CoA dehydrogenase deficiency, which leads to a build-up of the neurotoxin glutaric acid, causing GA1 (Goodman et al., 1998; Larson et al., 2019). GA1 can manifest in infancy or have a late onset, and can present with various neurological (e.g. neurodevelopmental impairment, epilepsy, dementia, tremor) and psychiatric (e.g. BPD, anxiety) symptoms (da Costa Ferreira et al., 2008; Goodman et al., 1998; Pokora et al., 2019; Ramsay et al., 2018; Sanju et al., 2020). Glutaric acid has also been identified as a biomarker for violent presentations in SSD (Chen et al., 2020). Although, to date, no direct associations between OCD and *GCDH*/GA1 have been established, it is of interest to note that late-onset GA1 can present with brain neoplasms and GA1 patients of any age may present with chronic kidney disease (CKD) (Afsoun Seddighi et al., 2015; Larson et al., 2019). A number of studies suggest that brain neoplasms and CKD are associated with higher risk for OCD or OCD-like symptoms (Afsoun Seddighi et al., 2015; Berthier et al., 1996; Chacko et al., 2000; Larson et al., 2019; Yousefichaijan et al., 2014; Yousefichaijan et al., 2016). Several studies have suggested that brain neoplasms and CKD can lead to central nervous system dysfunction, and higher prevalence of psychiatric and cognitive disorders (e.g. memory disorders, anxiety disorders, ADHD, OCD and MDD) (Arnold et al., 2016; Chaijan et al., 2015; Durand et al., 2018; Nur et al., 2019; Silva et al., 2019; Yousefichaijan et al., 2014). However, the exact causal relationships between brain neoplasms, CKD and OCD (and other neuropsychiatric phenotypes) remain unclear (Yousefichaijan et al., 2014; Yousefichaijan et al., 2016). The enrichment of *GCDH* pathogenic variants in OCD patients should be further explored to elucidate whether there is a true association and its pathobiological mechanism.

An additional OCD patient was found to have 2 LP variants within the *ASL* gene coding for the enzyme argininosuccinate lyase (ASL), biallelic mutations in which result in autosomal recessive argininosuccinic aciduria (Baruteau et al., 2017). ASL is one of the six enzymes involved in the breakdown and removal of nitrogen; consequently, the main presentation of argininosuccinic aciduria is hyperammonemia (Nagamani et al., 2012). Argininosuccinic aciduria can result in neurodevelopmental impairment, epilepsy, seizures and ADHD (Nagamani et al., 2012). To date, there has been one case study of a patient with refractory OCD and body dysmorphic disorder associated with hyperammonemia (Cleveland et al., 2009). Hyperammonemia has been strongly associated with various neuropsychiatric phenotypes, including dementia, psychosis, mood disorders and hallucinations (O. Bonnot et al., 2015; Enns et al., 2005; Leo et al., 2019). Significantly, antipsychotics and mood stabilizers used most commonly in the treatment of SSD and BPD are known to cause recurrent hyperammonemia in patients, resulting in encephalopathy and delirium (Muraleedharan et al., 2015; Y.-F. Wu, 2017). One study showed that up to one-third of SSD patients taking valproic acid were found to have hyperammonemia (Ando et al., 2017). Not surprisingly, administration of valproic acid and corticosteroids can worsen urea cycle disorders, such as argininosuccinic aciduria (O. Bonnot et al., 2015). This underlines the importance of accurate genetic diagnosis in psychiatric patients who may have an underlying genetic disorder for which standard psychotropic medications may be contraindicated. In particular, psychiatric patients presenting with hyperammonemia and/or worsening symptoms following the administration of certain antipsychotics or mood stabilizers should be prioritized for genetic evaluation for a possible IEM.

Genetic variants associated with non-IEM TGDs were also found to be enriched within the study cohort relative to the general population. The majority of these variants were found within ion channel genes associated with autosomal dominant disorders, with a total of 37 patients being putatively affected. Ion channel diseases, known as channelopathies, have been increasingly studied in the field of neuropsychiatry, given that the brain is an electrically excitable tissue (Gargus, 2006; Schmunk et al., 2013). Channelopathies result from mutations in calcium, sodium, potassium and/or chloride ion channels, all of which are involved in a wide array of physiological processes, including neurotransmission, secretion and cell proliferation, among others (Imbrici et al., 2016). Channelopathies affecting the central nervous system include cerebellar ataxia syndromes, epileptic encephalopathies and familial hemiplegic migraine (Imbrici et al., 2016). Interestingly, channelopathy phenotypes such as epilepsy are also comorbid for other neurodevelopmental disorders, including autism spectrum disorder (ASD), intellectual disability, parkinsonism, SSD, BPD and others (Fanella et al., 2020; Imbrici et al., 2016; Knott et al., 2016). Most importantly, channelopathies can respond to specific medications that modulate ion channel activity, such as acetazolamide or valproate (Camia et al., 2017; Chiron et al., 2011; Cleland et al., 2008; Imbrici et al., 2016). Novel therapies, including gene therapy, gene editing and animal toxins, are also being explored for the treatment of channelopathies (Kozlov, 2018; Wykes et al., 2018).

One of the genes with the most variants found within the study sample (n = 20) was *CACNA1A*, which encodes the calcium voltage-gated channel subunit alpha-1 (Indelicato et al., 2019). Mutations in *CACNA1A* result in dominantly inherited familial hemiplegic migraine, epileptic encephalopathy, episodic ataxia type 2 and/or spinocerebellar ataxia type 6 phenotypes (Indelicato et al., 2019). Furthermore, *CACNA1A* variants have been associated with ASD, SSD and BPD in numerous psychiatric genetic studies (Damaj et al., 2015; Indelicato et al., 2019; Li et al., 2017; McCarthy et al., 2016). Interestingly, variants in *CACNA1A* have been shown to play a role in antipsychotic treatment response in SSD patients (O’Connell et al., 2019). There were also numerous variants (n = 18) identified within the voltage-gated sodium channel genes, *SCN1A, SCN2A* and *SCN3A*. Mutations in the *SCN* genes are associated with autosomal dominant generalized epilepsy with febrile seizures, Dravet syndrome and/or familial hemiplegic migraine (Miller et al., 2019; Wolff et al., 2017; Zaman et al., 2020). There is also increasing evidence for the association of *SCN1A, SCN2A* and *SCN3A* variants with ASD, SSD and BPD, with genotype-phenotype correlations showing that the functional effect of a given mutation tends to be associated with a specific set of clinical presentations (Bartnik et al., 2011; Carroll et al., 2016; Nickel et al., 2018; Suddaby et al., 2019; Yamakawa, 2016).. For example, *SCN2A* loss-of-function mutations are generally associated with neurological phenotypes (e.g. ataxia, epilepsy), while gain-of-function mutations tend to be associated with neuropsychiatric phenotypes (e.g. ASD, intellectual disability), and some variants with both gain- and loss-of-function effects can result in a broader phenotypic spectrum (Suddaby et al., 2019; Winquist et al., 2018). Furthermore, using gene set-based analytics testing on genome-wide association study data, Askland *et al*. (2012) found ion channel genes to be consistently enriched in SSD samples across both European-American and African-American ethnic groups, suggesting that variations within ion channel genes could play a role in SSD genetic susceptibility (Askland et al., 2012). In this study, 37 patients were identified to be putatively affected with a channelopathy (one patient had two different *CACNA1A* variants), with one-third (n = 12) diagnosed with SSD, mirroring the known strong association of ion channel dysfunction and SSD susceptibility. Of the SSD patients, four patients were identified to have *CACN1A1* pathogenic/LP variants. Although there was no direct measure of treatment-responsiveness in these patients, 3 (25%) had reportedly been treated with clozapine or olanzapine, which are often second-line antipsychotics used after incomplete response to first-line antipsychotics. This provides further support for the role of voltage-gated calcium channel gene variants in SSD treatment response (O’Connell et al., 2019).

Of interest, another third (n = 11) of the putative channelopathy patients were diagnosed with OCD. Although there is no direct association known between *CACNA1A* and *SCN* gene variants, and OCD, there is some literature suggesting a potential link between ion channel genes and OCD-like symptoms/behaviours. Specifically, familial hemiplegic migraine patients and mouse models with *CACNA1A* mutations have been reported to present with comorbid OCD or OCD-like behaviours (Bøttger et al., 2012; Bøttger et al., 2016; Dehghani et al., 2019; Marconi et al., 2003; Santoro et al., 2011). Interestingly, calcium channel antagonists showed inhibition of OCD behaviours in OCD mouse models and had anxiolytic effects (Bandelow, 2008; Egashira et al., 2008), while sodium channel activators have been shown to induce anxiety- and OCD-like behaviours in rats (Saitoh et al., 2015). These studies support a potential shared molecular pathophysiology between ion channel disorders and OCD-related phenotypes.

Two patients were found to be putatively affected with a channelopathy, as well as another genetic disease. Specifically, one male OCD patient was found to have pathogenic variants in both *SCN1A* and *OTC*, while a female SSD patient was found to have pathogenic variants in *SCN1A* and *PDHA1*. Interestingly, OTC deficiency patients can present with cerebellar ataxia, seizures and epilepsy, all of which are phenotypes also prevalent amongst channelopathies (Barkovich et al., 2020; Crowe et al., 2018; Hidaka et al., 2020; Im et al., 2018; Pizzi et al., 2019; B. Wu et al., 2018). Moreover, OTC deficiency is considered a differential diagnosis for hemiplegic migraine (Kumar et al., 2020). Similarly, PDHA1 deficiency patients can also present with neurological phenotypes akin to channelopathies (Bhandary et al., 2015; Debray et al., 2008; Prasad et al., 2011). Even more striking is the finding of anti-PDHA1 antibodies in a subset of SSD patients (Nakagami et al., 2020). Nakagami *et al*. (2020) found that patients with anti-PDHA1 antibodies had increased volumes of the left occipital fusiform gyrus and left cuneus, a finding that is in contrast to the decreased volumes typically seen in SSD patients in comparison to controls (Nakagami et al., 2020). Increased volumes of fusiform gyrus have been previously associated with synesthesia – a disorder characterized by abnormal perception in response to the presence or absence of an external sensory stimulus – which can potentially be linked to an underlying common mechanism for hallucinations and delusions typical of SSD (Bouvet et al., 2017; Hupé et al., 2015; Weiss et al., 2009). PDHA1 dysfunction has been associated with mitochondrial dysfunction, as well as numerous neuropsychiatric disorders, including Parkinson’s disease, dementia, psychosis and SSD (Nakagami et al., 2020). A patient with *PDHA1* mutation was diagnosed with SSD after presenting with auditory hallucinations, psychosis and seizures, suggesting the potential for a subgroup of SSD patients to be influenced by genetic disorders that can cause organic psychosis (Satogami et al., 2017). Most notably, PDHA1 deficiency patients have been shown to have favorable response to acetazolamide, a drug commonly used to treat channelopathies (Egel et al., 2010; Livingstone et al., 1984; Platt et al., 2012).

Taken together, there is a growing body of evidence for shared pathogenetic mechanisms of channelopathies and psychiatric disorders, with important implications for development of targeted therapies that can potentially be effective across the various brain phenotypes associated with ion channel genes (Gargus, 2006; Imbrici et al., 2016). This further emphasizes the need for accurate genetic diagnosis of psychiatric patients so that precision therapies can be initiated.

A major limitation of our study was the paucity of clinical phenotypic data beyond the psychiatric diagnoses and medications taken at the time of recruitment. Statistical analysis of this data did not reveal any differences in the average number of medications for patients with and without genetic disease variants (data not shown). We were not able to postulate based on the available clinical data whether patients had non-psychiatric findings consistent with the genetic disorders identified in this study. However, all patients included in this study consented to be re-contacted, enabling future follow-up to clinically validate the study findings. Another limitation of the study analysis was the assumption of HWE in calculating carrier frequencies. Comparing the established rate of female heterozygous *ABCD1* mutation carriers at 9.52E-06 to the calculated rate of 4.5E-03 based on HWE, it can be seen that the calculated rate is significantly higher than the observed heterozygote rate (Wiesinger et al., 2015). Similarly, the calculation of rates of females heterozygous for *OTC* mutations and carrier rates for recessive disorders where established carrier rates were not known/available based on HWE could have led to an inflated estimation. This may contribute to the apparent lack of enrichment of variants in these genes in analysis of the study results. Finally, future replication studies should include carefully clinically screened control samples for direct statistical comparison purposes.

In conclusion, this is the first direct, comprehensive study of the burden of TGDs in a large, varied psychiatric cohort. The results of this study support that pathogenic gene variants associated with TGDs are enriched in primary psychiatric populations. This strongly suggests that the prevalence of these, and most likely many other, genetic diseases are greatly underestimated in psychiatric populations. Increasing awareness and ensuring accurate diagnosis of TGDs will open new avenues to targeted treatment for a subset of psychiatric patients. Future studies into the psychiatric sub-phenotypes associated with TGDs and whether carriers of these disorders may exhibit clinical manifestations, as well as the response of psychiatric symptomatology to targeted treatments will be vital in optimizing the precision diagnosis and management of many psychiatric patients.

## 1.6 Acknowledgements

The authors would like to thank Dr. John B. Vincent, Dr. Clement Zai, and Anna Mikhailov for their technical assistance, and critical and constructive input during this study.

## 1.7 Conflict of Interest

All authors declare no conflict of interest.

## 1.8 Funding Information

V.S. was funded by the Canadian Institute of Health Research (CIHR) doctoral award. This project was funded by an unrestricted research grant from Recordati Rare Diseases.

## Notes

### Competing Interest Statement

The authors have declared no competing interest.

